# A Common Recognition Architecture Across Memory Scales

**DOI:** 10.64898/2026.11.04.764437

**Authors:** Chong Zhao, Audrey Kim, Sintra Horwitz, Temilade Adekoya, Edward K. Vogel

## Abstract

Recognition decisions occur across memory episodes that vary vastly in scale, from compact single-item episodes to extended experiences involving hundreds of items. To test how memory scale shapes the neural mechanisms underlying old/new judgments, we recorded EEG while participants performed a recognition memory task across set sizes ranging from 1 to 128 items (spanning 2 to 256 recognition judgments). Classic electrophysiological signatures of recognition, including frontal and parietal old/new effects, persisted continuously across set sizes. Furthermore, multivariate neural classifiers trained on small memory scales generalized robustly across set sizes. Rather than revealing distinct neural formats, increasing memory scale produced systematic, continuous shifts in the timing and strength of a shared recognition response: signals emerged progressively later and weaker at larger memory scales. Temporal-alignment and cross-correlation analyses demonstrated that responses across scales followed a common waveform trajectory, with timing shifts propagating into motor response preparation. Finally, individual difference analyses revealed that this common recognition format exhibits scale-dependent behavioral relevance, tracking working-memory capacity at smaller scales and long-term memory performance at larger scales. Together, these findings demonstrate that recognition signals share a common neural format across scale while changing dynamically in timing, strength, and behavioral relevance, providing evidence for convergence onto a shared recognition architecture.

## Introduction

A central function of memory is to allow past experience to guide present decisions. In recognition, this requires determining whether a current stimulus has been encountered before. Such decisions can operate across very different scales. At the smallest scale, we may judge whether a single object was seen moments earlier; at larger scales, the same old/new decision may require access to many objects encoded across an extended experience and queried throughout a prolonged sequence of test decisions. These situations differ not only in the number and accessibility of the relevant memories, but also in the temporal extent of encoding and the duration over which those memories must support recognition judgments. We use the term memory scale to capture this combined informational and temporal extent of a memory episode.

For more than a century, theories of memory have sought to explain how information remains immediately accessible in some circumstances but must be recovered from a more enduring store in others. William James (James, 1890) characterized this distinction as primary and secondary memory, a framework later formalized in the modal model of memory as separate short-term and long-term memory stores (Atkinson & Shiffrin, 1968). Modern theories similarly distinguish working memory (WM), a capacity-limited system that supports the temporary maintenance and manipulation of information, from long-term memory (LTM), which supports the storage of vast quantities of information over extended periods. One of the most striking differences between these systems concerns capacity. Visual working memory is typically limited to approximately three to four objects (Cowan, 2001), whereas visual long-term memory can store thousands of objects with remarkable fidelity and detail (Brady et al., 2008; Wolfe et al., 2023). Viewed through the lens of memory scale, this traditional distinction contrasts compact episodes involving a few highly accessible representations with larger and more temporally extended episodes involving many less accessible representations.

Differences in capacity do not necessarily imply that WM and LTM are separate storage systems. State-based accounts propose that WM reflects a privileged activation state of representations that ultimately reside within long-term memory. In Cowan’s embedded-processes model, for example, working memory consists of a subset of long-term memory representations that have been temporarily activated, with an even smaller subset occupying the focus of attention (Cowan, 1999, 2001). Related frameworks similarly emphasize differences in activation, accessibility, and attentional prioritization rather than distinct storage systems (Oberauer, 2002). From this perspective, changes in memory scale may alter the accessibility of relevant representations without necessarily changing the fundamental system though which they are stored or evaluated.

Although these theoretical perspectives differ substantially in their assumptions regarding memory storage, they share a common challenge: explaining how stored information is ultimately used to make recognition decisions. Whether a memory representation resides in a dedicated working memory store, an activated portion of long-term memory, or a currently attended subset of activated representations, successful recognition requires transforming stored information into a judgment about whether a stimulus has been encountered before. Theories that distinguish between memory stores and those that emphasize differences in activation therefore leave open the same question: Does the process that evaluated memory evidence remain stable as the scale of the memory episode expands? Surprisingly, relatively little is known about whether the neural processes supporting such recognition remain continuous across memory scales or change as relevant representations become more numerous, less accessible, and distributed across longer periods of time.

Previous research has largely focused on relationships between WM and LTM at encoding. Individual differences in WM capacity predict subsequent long-term memory formation (Fukuda & Vogel, 2019), and measures of WM and LTM ability are moderately correlated across individuals(Unsworth et al., 2014; Zhao & Vogel, 2025).

These findings suggest that the two systems share important constraints, potentially arising from common attentional or encoding bottlenecks. Far less is known, however, about the relationship between WM and LTM at retrieval, particularly whether the neural processes that transform stored information into recognition decisions remain consistent as memory scale increases.

At least three possibilities can be envisioned. The first is a **separate-retrieval** account. According to this view, information maintained within WM remains directly accessible and is evaluated through retrieval mechanisms distinct from those used to recover information from LTM. Recognition judgments based on WM and LTM should therefore recruit qualitatively different neural processes. Viewed across memory scale, this account predicts a discontinuity: recognition should change qualitatively as memory episodes expand beyond a few highly accessible representations to larger, temporally extended sets of candidate memories.

A second possibility is a **convergence** account. Under this framework, WM and LTM may rely on different retrieval operations, but the resulting evidence converges onto a common recognition architecture. Information may originate from distinct sources depending on its accessibility, yet once memory evidence becomes available, it is evaluated through shared neural computations that support old/new decisions. This account predicts that neural signatures associated with recognition should generalize across memory scales despite differences in the accessibility of the underlying memory representations. It also shows the behavioral relevance of a shared recognition signal to vary with scale: the same neural format may reflect evidence supplied by highly accessible representations at smaller scales and by the retrieval of less accessible representations at larger scales.

A third possibility is a **shared-access** account. Consistent with several activation-state theories, this perspective proposes that WM and LTM differ primarily in activation state rather than retrieval mechanism. Recognition judgments would therefore be expected to rely on highly similar neural computations across memory scales, with differences emerging primarily in the speed or strength with which information becomes available. Under this account, recognition processes should exhibit substantial continuity across conditions traditionally associated with WM and those requiring LTM retrieval. In its strongest form, the shared-access account predicts not only a common neural format across neural scales, but also that individual differences in recognition-related activity at smaller and larger scales should reflect a common underlying ability.

Distinguishing among these possibilities has proven difficult because the neural study of recognition memory has largely evolved along two separate tracks. One tradition has focused on long-term memory retrieval and has identified robust electrophysiological signatures of recognition, including the frontal and parietal old/new effects (Rugg & Curran, 2007). These signals reliably distinguish previously encountered items from novel items and have frequently been linked to familiarity- and recollection-related processes. A second, smaller literature has documented similar old/new effects in short-term recognition tasks. For example, Danker and colleagues (Danker et al., 2008) reported both FN400 and LPC old/new effects during a Sternberg memory task and found that these components exhibited many of the same functional properties previously described in long-term memory paradigms. These findings have led several authors to suggest that common recognition processes may contribute to both short- and long-term memory judgments. However, because short-term and long-term recognition have typically been examined in separate paradigms, using different stimuli, task structures, and theoretical assumptions, direct comparisons have remained difficult. Consequently, it remains unclear whether similar old/new effects reflect a shared recognition process operating across memory scales, distinct access processes that converge on a common decision stage, or merely superficial similarities in scalp-level activity.

The present study was designed to bridge these traditionally separate literatures by examining recognition memory across a continuous range of memory scales within a single paradigm. We operationalized memory scale using set sizes ranging from 1 to 128 items. Importantly, this manipulation varied more that item number alone. As set size increased, the memory episode also expanded in study duration, study-test lag, accumulated interference, and the length of the subsequent recognition sequence. The smallest condition consisted of repeated, compact single-item study-test episodes, whereas the largest condition required subjects to encode 128 objects and then make 256 recognition judgements about that extended episode. We therefore use set size to refer to the specific experimental manipulation and memory scale to refer to the broader informational and temporal demands that varied with it.

At the smallest memory scales, recognition could plausibly rely on the highly accessible representations typically associated with WM. At the largest scales, successful recognition places much greater demands on access to less active representations distributed across an extended memory episode. Rather than assuming that these conditions map categorically onto WM and LTM, we treat them as points along a continuum of memory scale. Critically, the task and the old/new decision remained identical across all conditions, allowing us to ask whether the neural processes supporting recognition changed qualitatively or remained continuous as the informational and temporal demands of the episode increased.

We first examined two classic electrophysiological signatures of recognition memory, the frontal and parietal old/new effects, to determine whether they emerged continuously across memory scales. We then used multivariate decoding to test whether neural representations supporting old/new discrimination generalized across set sizes. Together, these analyses provide a test of the **separate-retrieval** account: if WM- and LTM-based recognition depend on qualitatively distinct retrieval processes, recognition-related neural activity should differ sharply across memory scales and classifiers should show limited cross-scale generalization. In contrast, both the convergence and shared-access accounts predict continuity in recognition-related activity and generalization of neural representations across memory scales.

We next tested whether increasing memory scale shifted a common recognition response in time and strength. If scale primarily affects the accessibility of memory evidence, larger set sizes should not simply abolish or replace the recognition signal. Instead, they should delay and weaken a similar response. We tested this prediction by examining the joint relationship between old/new amplitude and latency and by asking whether these old/new waveforms across set sizes could be aligned through temporal-lag analyses. These analyses allowed us to determine whether differences across memory scales reflected distinct recognition processes or temporally shifted expressions of a common response.

Finally, we tested whether individual differences could distinguish between these latter two accounts. The **convergence** account allows recognition-related neural activity to be shared in overall form while retaining some memory-system specificity within individuals. By contrast, the strongest form of the **shared-access** account predicts that the neural activity supporting retrieval at smaller memory scales should also predict performance at larger memory scales, and vice versa, because both would reflect the same underlying access mechanism. Thus, the present study asks two linked questions: first, whether recognition-related neural activity generalizes across memory scales; and second, whether that shared activity is best characterized as convergence onto a common recognition architecture or as evidence for a fully shared access mechanism. We therefore examined whether recognition-related neural activity at smaller set sizes was associated with independent measures of working memory ability, whereas activity at larger set sizes was associated with long-term recognition performance. This analysis tested whether a common neural recognition format draws on different sources of memory evidence as scale increases.

By examining recognition from compact single-item episodes to extended episodes involving 128 studied objects and 256 recognition judgments, the present study asks how the brain supports old/new decisions as memory episodes expand in informational and temporal scale. A separate-retrieval account predicts discontinuities in recognition-related neural activity across set size. In contrast, the convergence and shared-access accounts predict that increasing memory scale will primarily alter when and how strongly a common recognition signal emerges. These accounts differ, however, in whether that shared signal should exhibit the same behavioral relationships across scales. Together, the present analyses test whether recognition relies on distinct processes at different memory scales, a common access mechanism, or different sources of memory evidence that converge on a shared recognition architecture.

## Method

### Participants

One hundred and eleven participants took part in the study (mean age = 25.1 years). Sample size was determined using G*Power, aiming to achieve 80% power to detect a small effect size (r = 0.3) at the conventional alpha level of .05. All participants reported normal or corrected-to-normal vision and normal color perception, and no history of neurological disorder. Study procedures were approved by the relevant University of Chicago Institutional Review Board and participants were compensated for their participation.

### Stimuli and Procedure

Images were selected from the THINGS dataset (Hebart et al., 2023), a database with 1854 unique concepts and 26107 images. One exemplar was selected from each concept so that no repeated concept was used throughout each session (i.e., apple versus avocado). The recognition memory procedure for all set sizes (1, 2, 4, 8, 16, 32 and 128) followed established protocols in the literature and consisted of separate study and test phases (Fukuda & Woodman, 2015; Zhao & Woodman, 2020). The experiment included set sizes of 1, 2, 4, 8, 16, 32, and 128. Each encoding condition consisted of a total of 128 trials. This was achieved by varying the number of blocks across set sizes, such that participants completed 128 blocks in the set size 1 condition, with progressively fewer blocks as set size increased, down to a single block in the set size 128 condition (see **Fig. 1**).

**Fig 1.**
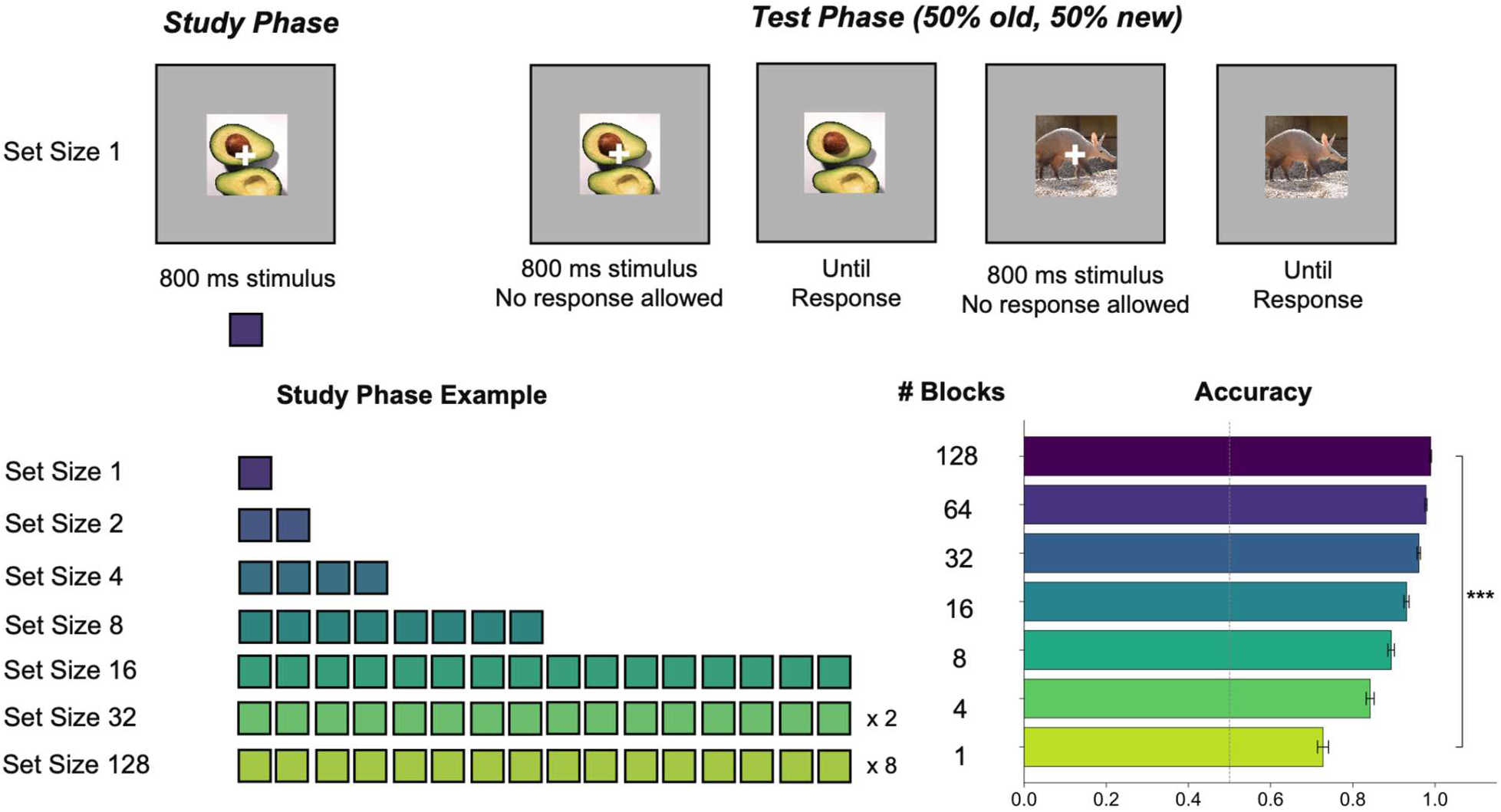
Visual Recognition Memory Paradigm with varying memory scales (set size 1, 2, 4, 8, 16, 32 and 128).

In the set size 1 condition, participants viewed a single image presented at the center of the screen for 800 ms during the encoding phase. Stimuli were displayed against a gray background (90.0 cd/m²), and a white fixation cross was superimposed on the image to encourage central fixation and discourage eye movements. The interstimulus interval was randomly jittered between 250 and 400 ms.

Following the encoding phase, participants completed a recognition test trial in which a single image was displayed at the center of the screen for 800 ms with a central white fixation cross. To minimize motor and ocular artifacts, participants were instructed to remain still while the fixation cross was visible. After 800 ms, the fixation cross disappeared and the image remained on screen until a response was made. Half of the test images had been presented during the encoding phase (old), and the remaining half were novel (new). The order of old and new images was randomized across trials. Participants indicated whether each image was “old” or “new” using designated keyboard keys (“z” for old; “/” for new).

All other set size conditions followed the same general structure, differing only in the number of images presented during the encoding phase and the corresponding number of test trials. Block order was randomized across participants.

### Preprocessing and artifact rejection of EEG signals

Participants were positioned in an electrically isolated booth, with their heads stabilized using a cushioned chinrest placed 74 cm from the display screen. Electroencephalographic (EEG) signals were recorded via 30 active silver/silver chloride (Ag/AgCl) electrodes integrated into a stretchable cap (actiCHamp system, Brain Products, Munich, Germany), arranged according to the international 10-20 placement standard (electrode sites: Fp1, Fp2, F7, F8, F3, F4, Fz, FC5, FC6, FC1, FC2, C3, C4, Cz, CP5, CP6, CP1, CP2, P7, P8, P3, P4, Pz, PO7, PO8, PO3, PO4, O1, O2, Oz).

Additional electrodes were adhered to the left and right mastoids using adhesive stickers, and a ground electrode was integrated at the Fpz site within the cap. EEG signals were initially referenced to the right mastoid and subsequently re-referenced offline to the average of both mastoids. The signal was bandpass filtered (0.01–80 Hz) with a 12 dB/octave roll-off and digitized at a sampling rate of 500 Hz. All electrode impedances were maintained below 10 kΩ.

To track ocular activity such as blinks and saccades, both electrooculographic (EOG) and eye-tracking data were recorded. EOG was captured using five passive Ag/AgCl electrodes: two for vertical EOG (above and below the right eye), two for horizontal EOG (approximately 1 cm lateral to each eye), and one ground electrode on the left cheek. Eye position was also monitored with a desktop-mounted EyeLink 1000 Plus system (SR Research, Ontario, Canada), operating at a sampling rate of 1,000 Hz.

### Artifact Rejection

For horizontal eye movements, we employed a sliding-window algorithm to identify horizontal eye movements using both horizontal electrooculogram (HEOG) signals and eye-tracking gaze data. For the HEOG-based detection, a split-half sliding-window method was applied with a 100 ms window moving in 10 ms increments. An eye movement was flagged if the voltage difference between the two halves of the window exceeded 20 µV. This HEOG-based artifact detection was used only in trials where eye-tracking data were unreliable or unavailable. In parallel, eye-tracking rejection was carried out by analyzing horizontal (x-axis) and vertical (y-axis) gaze positions using the same 100 ms window and 10 ms step size. A trial was excluded if eye position shifted more than 0.5° of visual angle within any given window.

To detect blinks, we applied a sliding-window analysis to the vertical EOG signal using an 80 ms window and a 10 ms step size. A blink was marked when the voltage change across the window exceeded 30 µV. Complementarily, we identified blink periods in the eye-tracking data by locating segments where positional data were missing, indicating that the eyes were closed.

### Parietal and Frontal Old/New Effect Analysis

To calculate the parietal and frontal old/new effects, we first identified sets of electrodes associated with each region. Based on prior literature, for the parietal old/new effect, we included electrodes PO3, PO4, PO7, PO8, P3, P4, P7, P8, and Pz. For the frontal old/new effect, we used electrodes F3, F4, F7, F8, and Fz. For each region, we computed the average event-related potential (ERP) time series across the selected electrodes to obtain a representative waveform. We then focused on correct trials within each set size and repetition condition and categorized them into two trial types: old (images that participants had previously studied) and new (novel images that had never been studied). We computed the average ERP for each trial type separately. The old/new effect was then derived by subtracting the ERP waveform for new trials from that for old trials, calculated separately for each participant. This difference waveform, named old/new effect, reflects neural activity specific to recognition memory.

## Results

To investigate how memory scale influences the neural processes supporting old/new judgments, we recorded scalp EEG while participants performed a recognition memory paradigm across set sizes ranging from 1 to 128 items. To examine how memory access changes across different memory scales, we recorded EEG while participants performed a recognition memory task. The critical manipulation was list set size: smaller memory scales, with list lengths within the typical range of visual memory capacity (approximately 4 items), were expected to place relatively greater demands on active maintenance, whereas larger memory scales, including supra-capacity lists of 32 and 128 items, were expected to place greater demands on retrieval from more durable memory representations.

A total of 111 participants encoded lists of images and subsequently judged whether each test image was old or new. Set size varied parametrically (1, 2, 4, 8, 16, 32, or 128 items; **Fig. 1**), sampling recognition across a continuum of memory scales ranging from repeated single-item episodes to an extended episode involving 128 studied objects and 256 recognition judgments. At test, half of the trials were previously encoded images and half were novel foils. To equate trial numbers across conditions, participants completed 128 blocks of set size 1, 64 blocks of set size 2, and so forth, down to 1 block of set size 128, yielding 128 encoding trials and 256 test trials per condition. Block order was randomized, and set size was not disclosed until completion of encoding. We used this paradigm to test whether increasing memory scale induces a qualitative shift in recognition processes, delays and attenuates a common decision signal, or alters the behavioral relevance of a shared neural architecture.

### Univariate Parietal old/new effect revealed convergent access to increasing memory scale during retrieval

Given that WM and LTM have vastly different properties with respect to capacity limits, activation levels, and even brain areas related to storage, we asked whether retrieval from smaller and larger memory scales rely on distinct access mechanisms, or whether they share a common neural process during memory-based decision-making. A key operation in such decisions is recollection, the recovery of an item together with details of its prior occurrence. Recollection has been studied extensively with larger memory scales (Rugg & Curran, 2007), but it remains unclear whether the same neural process supports retrieval when items fall within a much smaller memory scale. A key prediction of the **separate-retrieval** account is that recognition judgments based primarily on WM should recruit neural processes qualitatively different from those engaged when retrieval depends on LTM. In contrast, both the **convergence** and **shared-access** accounts predict continuity in recognition-related neural activity across memory scales.

We indexed recollection using the parietal old/new effect, a well-established univariate EEG signature characterized by more positive activity over parietal scalp sites for correctly recognized old items compared to new items. Under a **separate-retrieval** account, this signal should be prominent when retrieval depends on larger memory scale retrieval, but not further enhanced when decisions primarily rely on smaller memory scales, which may instead be supported by a distinct mechanism based on sustained neural activity. In contrast, a shared-access account predicts that the amplitude and timing of this signal should vary systematically with memory load, forming a continuous pattern across conditions that engage smaller and larger memory scales.

Across 111 participants, we found evidence against the **separate-retrieval** hypothesis, in that the parietal old/new effect was significant across all set sizes (1-128 items; all ps < 0.01, see **Fig. 2**). Thus, despite large differences in capacity limits and activation states, retrieval of items with varying memory scales recruited the same electrophysiological signature. Moreover, the amplitude of the parietal old/new effect decreases with increasing memory scales (F(6, 660) = 14.09, p < .001), suggesting that the memory strength decreases continually as memory scale increase. To further dissect this set size effect, we found that within the canonical WM capacity (1, 2, 4 items), amplitudes of parietal old/new effects did not differ from each other (p > 0.20). With increasing memory scales beyond WM capacity, however, the parietal old/new effect declined progressively, with marked reductions at set sizes 32 and 128. These findings rule out a strong **separate-retrieval** mechanism, with increasing memory scale associated with reduced memory strength.

**Fig 2.**
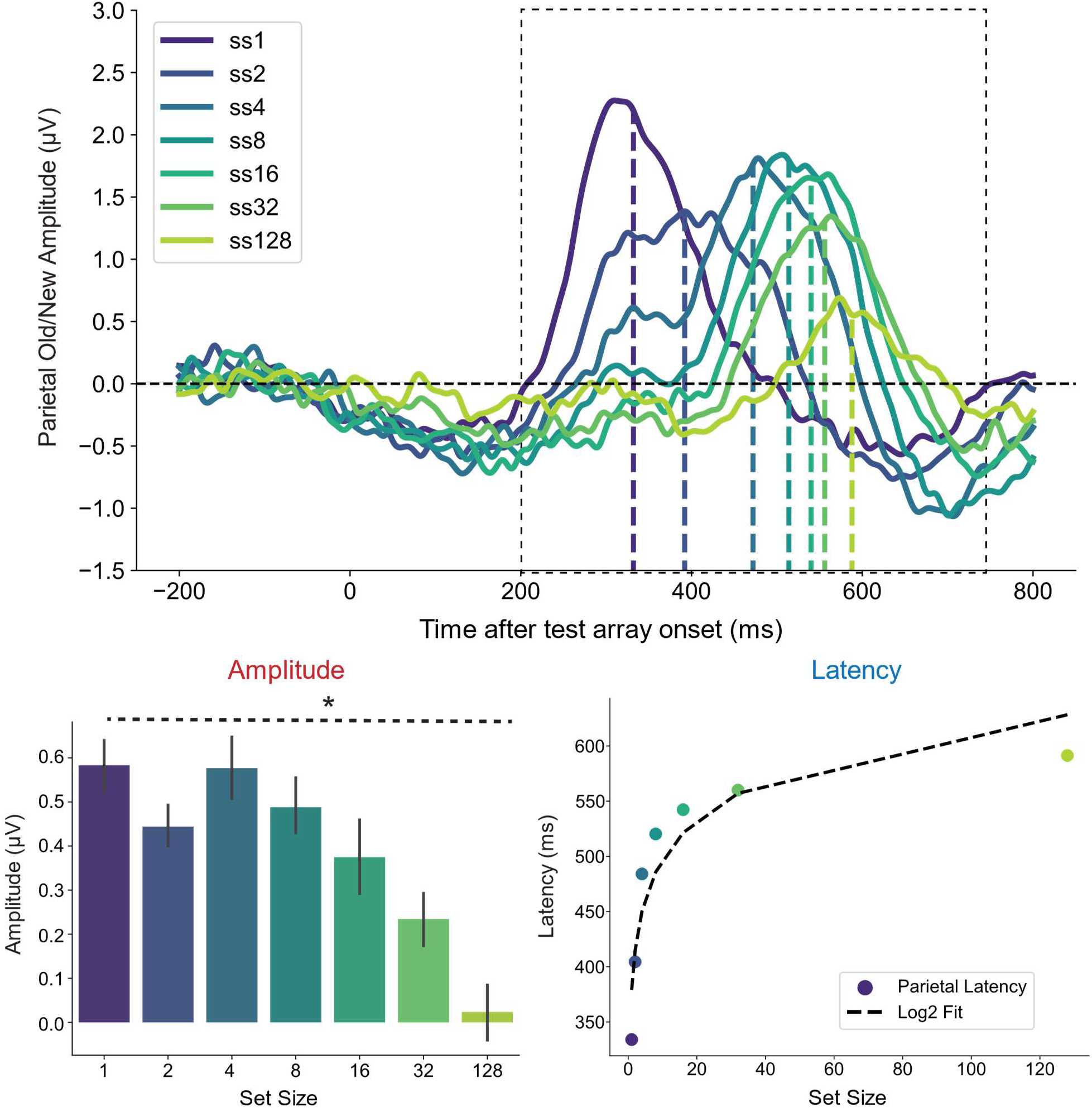
Amplitude and Latency of the Parietal old/new effect with varying memory scales (set size 1, 2, 4, 8, 16, 32 and 128).

Another critical factor in memory access is the speed with which items of interest are successfully retrieved during a task. A classic finding in the behavioral literature is that response times are faster when searching through a small set of memorized items than when searching through a larger set that exceeds smaller memory scales (Wolfe, 2012). If the shared-access hypothesis holds across both smaller and larger memory scales, the peak of the neural signature of recollection should mirror the response time patterns observed in memory search paradigms. To determine when the brain reaches peak evidence accumulation during memory-based decisions, we quantified the latency of the parietal old/new effect, defined as the time to reach 50% of the area under the curve (Kiesel et al., 2008). Supporting the shared-access account, peak latency occurred earlier for smaller set sizes and became progressively delayed as set size increased, with each set size differing significantly from adjacent set sizes. Interestingly, this increase in neural latency followed a logarithmic rather than linear function of set size. This logarithmic relationship suggests that although smaller and larger memory scales recruit a common access signal, their search dynamics differ. At smaller memory scales, adding a single item incurred a measurable temporal cost, consistent with a more serial search mechanism. In contrast, expanding the search space at larger memory scales (e.g., from 32 to 128 items) yielded diminishing latency costs, consistent with more parallelized retrieval. This logarithmic scaling mirrors prior behavioral evidence showing that response times in memory search increase as a log function of set size. Together, the amplitude and latency profiles of the parietal old/new effect support a shared or convergent access mechanism supporting retrieval across both smaller and larger memory scales.

### Univariate Frontal and parietal old/new effects persist across memory scale

We first tested whether classic electrophysiological signatures of recognition, the parietal and frontal old/new effects, persist continuously across memory scales or exhibit qualitative boundaries between smaller and larger set sizes. Our results suggest that the parietal old/new effect, a neural signature associated with recollection, emerges earlier and with greater amplitude with smaller memory scales, but is delayed and attenuated with larger memory scales. Despite these differences in temporal dynamics and magnitude, both forms of memory appear to rely on a common neural retrieval system at test. One plausible account is that recollection, defined as the binding of item and contextual information, constitutes a shared process across increasing memory scales. In contrast, familiarity-based processes may contribute more prominently to large-scale retrieval than to small-scale retrieval, given that shorter lists are more likely to rely on recollection. Under this **separate-retrieval** framework, one would predict that the neural correlate of familiarity, the frontal old/new effect, would be selectively engaged during retrieval of list with large memory scales and not exhibit earlier or stronger activation during retrieval of list with small memory scales.

Contrary to this prediction, our findings instead support either the **convergence** or **shared-access** account for familiarity-based recognition. Specifically, the frontal old/new effect was significant across all set sizes (1–128 items; all *p*s < .01, see **Fig. 3**), indicating that both small and large memory scale retrieval recruit a common electrophysiological signature of familiarity. Rather than disappearing or abruptly altering in topography at larger set sizes, both anterior and posterior recognition signatures persisted across the entire spectrum of memory scales.

**Fig 3.**
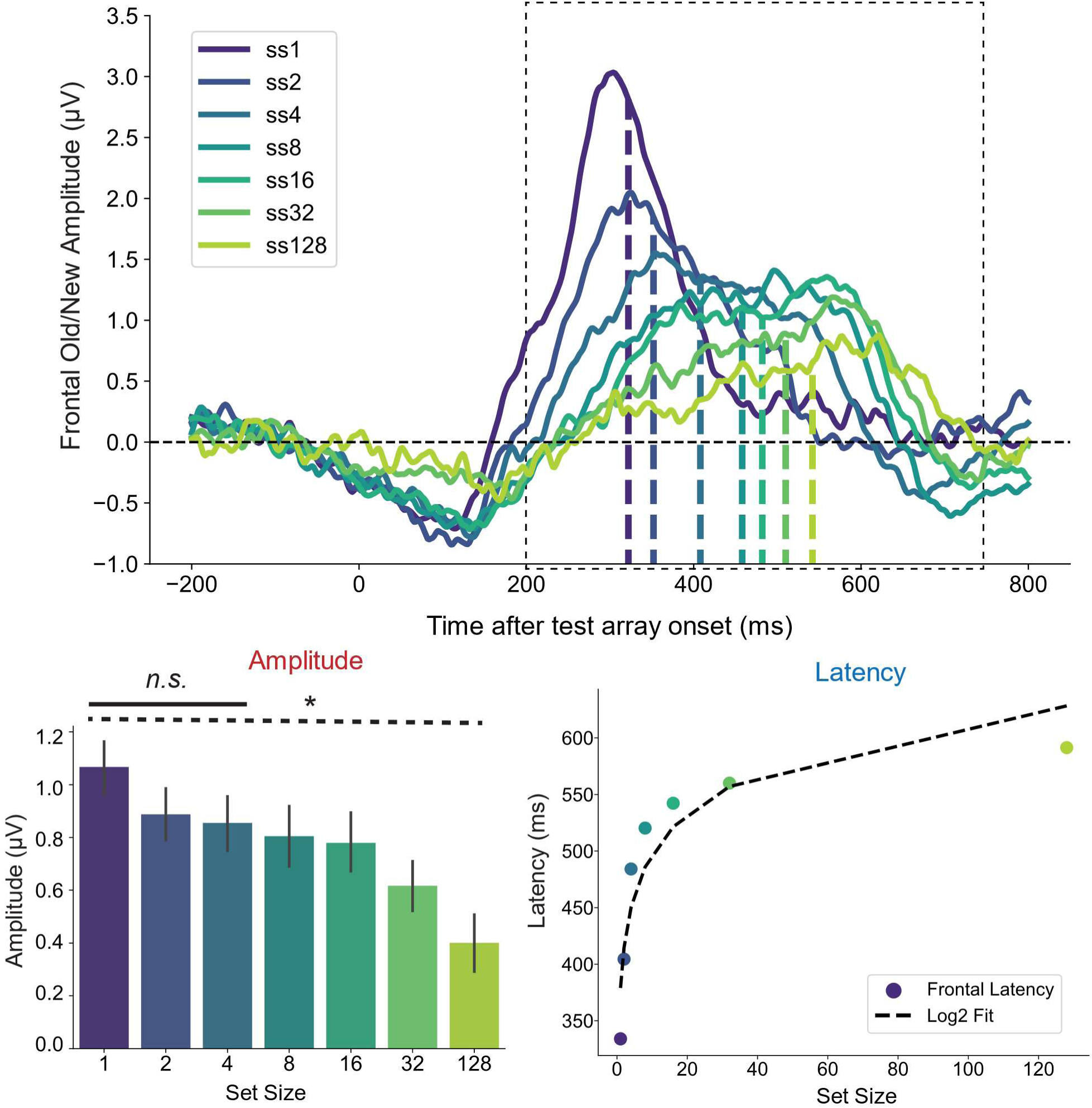
Amplitude and Latency of the Frontal old/new effect with varying memory scales (set size 1, 2, 4, 8, 16, 32 and 128).

In addition, the amplitude of the frontal old/new effect decreased systematically with increasing set size (F(6, 660) = 5.39, p < .001), suggesting a gradual reduction in the strength or accessibility of retrieved memory representations across memory scales. To further characterize this pattern, we compared set sizes within and beyond the range typically associated with smaller memory scales. Within the smaller-scale conditions (1, 2, and 4 items), frontal old/new amplitudes did not differ significantly (ps > 0.41). In contrast, at larger memory scales, the frontal old/new effect declined progressively, with particularly pronounced reductions at set sizes of 32 and 128 items. Together with the corresponding pattern observed for the parietal old/new effect, these findings indicate that recognition-related neural activity is relatively stable across smaller memory scales but becomes progressively weaker as memory scale increases. Rather than indicating a categorical distinction between memory systems, this graded pattern is consistent with a convergent or shared neural access mechanism in which increasing memory scale modulates the strength and accessibility of retrieved representations.

In addition to the amplitude finding, consistent with either the **convergence** or the **shared-access** hypothesis, peak latency occurred earlier for smaller set sizes and increased systematically with larger set sizes for frontal old/new effect. Notably, latency differences increased with set size (*p*s < .01), and this increase followed a logarithmic rather than a linear function of set size. This pattern again aligns with prior findings from memory search paradigms, which demonstrate that response time increases as a logarithmic function of memory set size.

### Temporal-alignment analyses show that responses across set sizes follow a common waveform trajectory

Lastly, we asked whether the parietal and frontal old/new effects reflect a coherent neural representation of memory across different memory scales. We examined the cross-correlation of the old/new effects across set sizes and recorded the peak cross-correlation across lags relative to set size 1. A high cross-correlation suggests that the neural signals at larger set sizes share a similar representational structure with those at set size 1, differing primarily by a temporal lag. For both parietal (**Fig. 4A**) and frontal (**Fig. 4B**) old/new effects across set sizes, we observed extremely high peak cross-correlations (rs > 0.89, significantly higher than mean cross correlation strength across all lags, ps < 0.001), indicating that memory scale does not substantially alter the neural representation of recognition memory despite substantial differences in set size.

**Fig 4.**
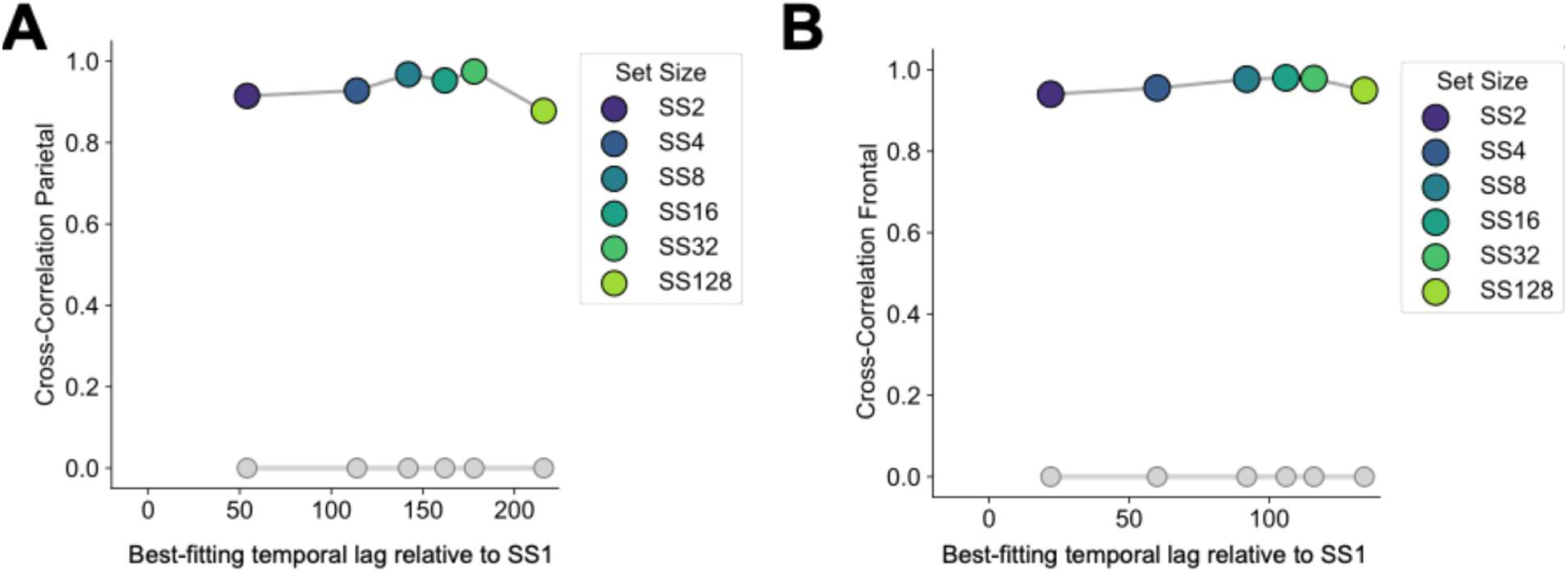
High peak cross-correlation of parietal (A) and frontal (B) old/new effects across set sizes, indicating that memory scale does not substantially modulate neural representations of recognition memory despite of vastly different memory scales.

Taken together, the findings from both the parietal and frontal old/new effects indicate that the amplitude and latency characteristics of these signals provide converging evidence for either a **convergent** or a **shared-access** mechanism underlying retrieval from across memory scales, while also revealing systematic differences in search dynamics across memory systems.

### Recognition-related neural patterns generalize across set sizes

Our univariate results collectively suggest that recognition may rely on a common neural access mechanism across different memory scales. However, because these univariate signatures were derived primarily from prior research with large memory scale, they may be influenced by biases in the existing literature and by assumptions about which neural features are relevant for recognition. To address this question more directly, we used a multivariate classification approach to discriminate old from new trials at set size 4, which served as the template condition. Set size 4 was selected because it represents a relatively small memory scale near the upper range of commonly observed working-memory capacity. The classifier therefore learned to distinguish “old” from “new” items based on distributed patterns of neural activity associated with memory access under relatively constrained conditions. Critically, if the resulting decision boundary captures a general neural operation supporting recognition, rather than features specific to the set size 4 condition, it should generalize across memory scales. Successful generalization to both smaller and larger memory scales would indicate that the neural code supporting old/new discrimination is shared across conditions, consistent with a convergent memory-access account. In contrast, limited or absent generalization would suggest that recognition relies on partially distinct neural processes as memory scale changes.

The classifier was trained on the test phase of the set size 4 condition. As shown in **Fig. 5**, multivariate decoding generalized robustly across set sizes, with significant temporal clusters indicated by the white dashed box. Notably, for set sizes 1 and 2, decoding of old versus new trials preceded that observed in the set size 4 training condition, suggesting faster signal discrimination for smaller memory loads, consistent with our univariate latency findings. In contrast, as set size increased beyond four items, decoding latencies lagged relative to the set size 4 template, indicating slower retrieval dynamics.

**Fig 5.**
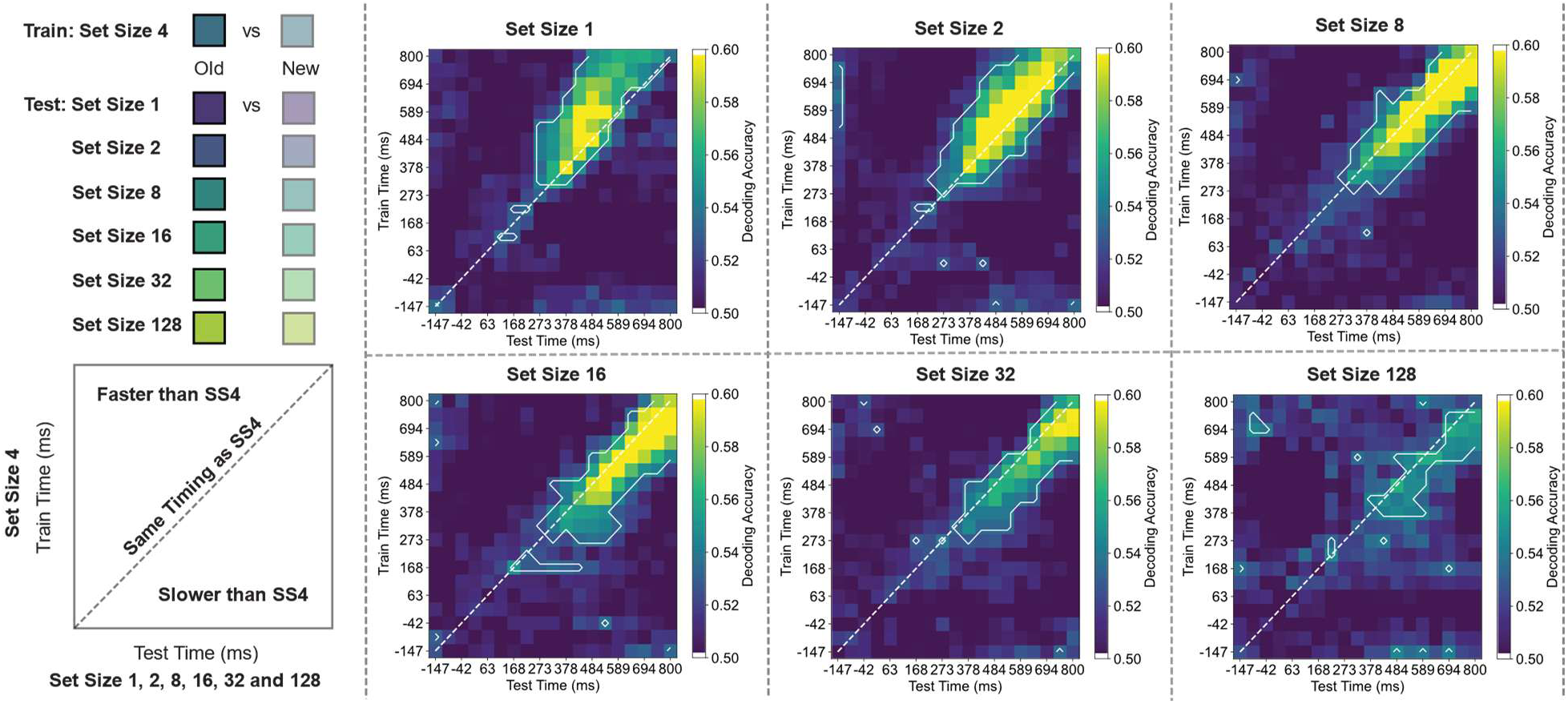
Multivariate Decoding of old/new items using single-trial EEG with varying memory scales (set size 1, 2, 4, 8, 16, 32 and 128).

Together, these findings suggest that although memory search operates via a shared mechanism across varying memory scales, retrieval speed is modulated by set size. Specifically, searching through larger memory sets is slower than accessing items actively maintained within smaller memory sets. This pattern indicates that the fine-grained spatio-temporal neural patterns that distinguish old from new items share a common neural format across smaller and larger memory scales.

### A common recognition signal shows scale-dependent behavioral relationships

Thus far, our analyses have focused on within-subject evidence to test whether recognition relies on a common neural access mechanism across memory scales.

However, individuals vary substantially in memory ability, and performance at smaller and larger memory scales may also be correlated. This raises a complementary question: in a well-powered sample, do neural measures of memory access predict individual differences in memory performance across scales? Critically, individual differences provide a way to distinguish between the **convergence** and **shared-access** accounts. The **convergence** account predicts that neural signatures may be shared across smaller and larger memory scales while retaining some degree of scale-specificity, such that neural measures associated with one scale need not predict individual differences in performance at another. In contrast, the strongest version of the **shared-access** account predicts that individual differences in neural activity during memory retrieval at smaller scales should predict individual differences in performance at larger scales, and vice versa, because both rely on the same underlying neural access mechanism. We therefore tested these competing predictions by examining whether individual differences in neural measures of retrieval at smaller memory scales predicted individual differences in performance at larger scales, and whether the reverse relationship was similarly observed.

Of the 111 participants, 82 completed an independent WM task with smaller memory scale (the change localization task), which provides a sensitive and reliable measure of WM capacity. We found that the parietal old/new effect for shorter lists significantly predicted WM performance (set sizes 1-32: rs = 0.23 -0.33, FDR-corrected ps < .05; see **Fig. 6**), but not for the largest set size (set size 128; r = 0.14, p = .22, BF = 0.4). Because these correlations may partly reflect LTM contributions, we next computed partial correlations controlling for LTM performance (indexed by set size 128 accuracy). The relationship between the parietal old/new effect and WM remained significant for smaller set sizes (set sizes 1-4: rs = 0.22-0.27, ps < .05), was mixed at intermediate set sizes (set size 8: r = 0.22, p = .05; set size 16: r = 0.27, p = .01), and was no longer reliable for larger set sizes (set size 32: r = 0.17, p = .13; set size 128: r = 0.04, p = .74). Moreover, frontal old/new effect did not predict any of the WM ability measures (ps > 0.30). This pattern suggests that the parietal old/new effect for smaller set sizes primarily tracks individual differences in WM ability.

**Fig 6.**
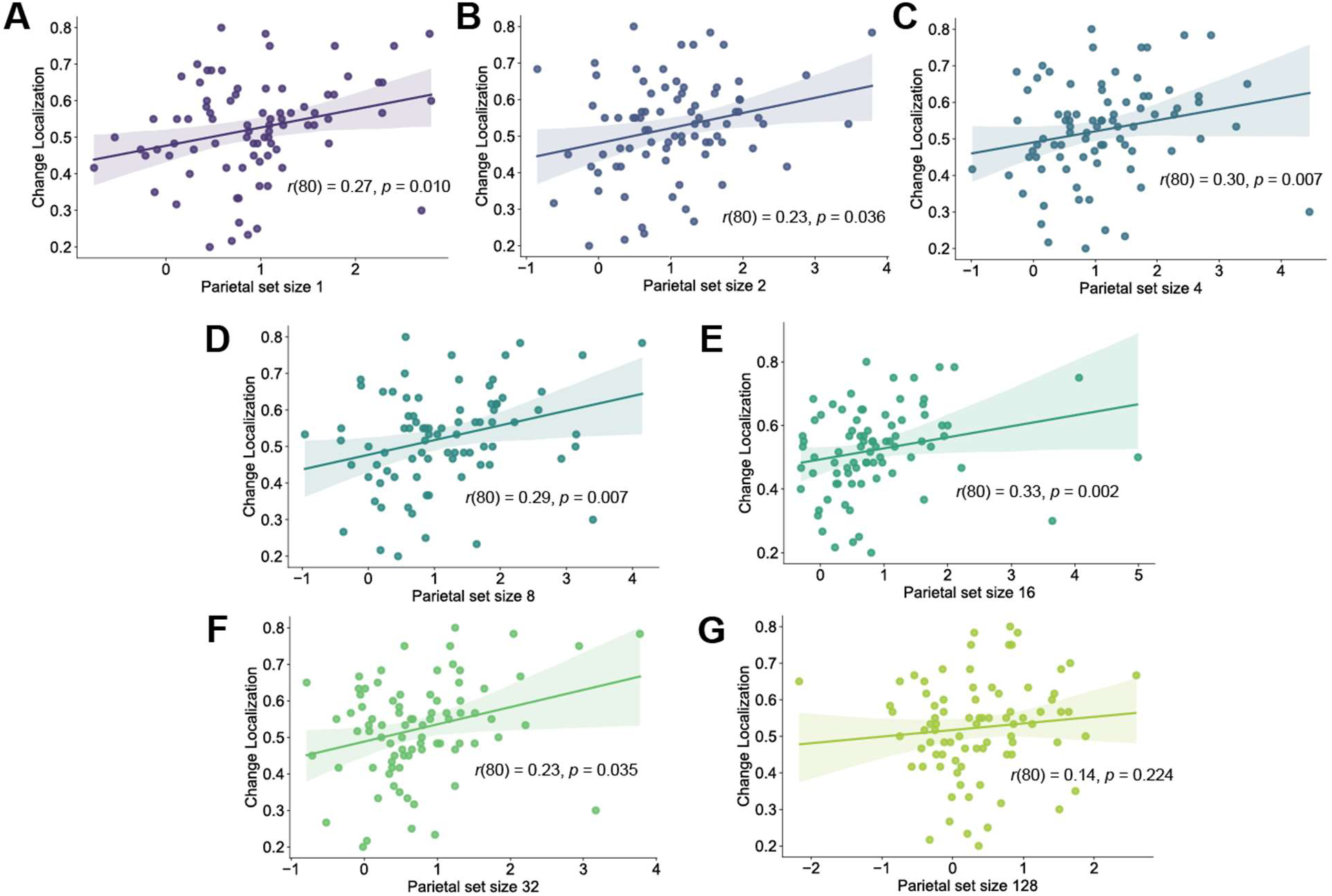
Parietal old/new effect for smaller set sizes selectively predict working memory performance.

We next asked whether the same neural measure predicts LTM ability with large memory scale. Correlational analyses revealed that the parietal old/new effect for very small set sizes (1–2) did not predict LTM performance (set size 128 accuracy; set size 1: r = 0.05, p = .63, BF = 0.2; set size 2: r = 0.03, p = .80, BF = 0.16), whereas effects emerging at and beyond WM capacity (set sizes 4-128) were significantly associated with LTM performance (rs = 0.24-0.33, FDR-corrected ps < .05; see **Fig. 7**). After controlling for WM ability, this relationship remained absent for small set sizes (set size 1: r = -0.03, p = .77; set size 2: r = -0.04, p = .73; set size 4: r = 0.17, p = .13), was mixed at intermediate set sizes (set size 8: r = 0.26, p = .02; set size 16: r = 0.21, p = .06; set size 32: r = 0.19, p = .09), and remained robust for the largest set size (128: r = 0.31, p < .01). Moreover, frontal old/new effect did not predict any of the LTM ability measures (ps > 0.16).

**Fig 7.**
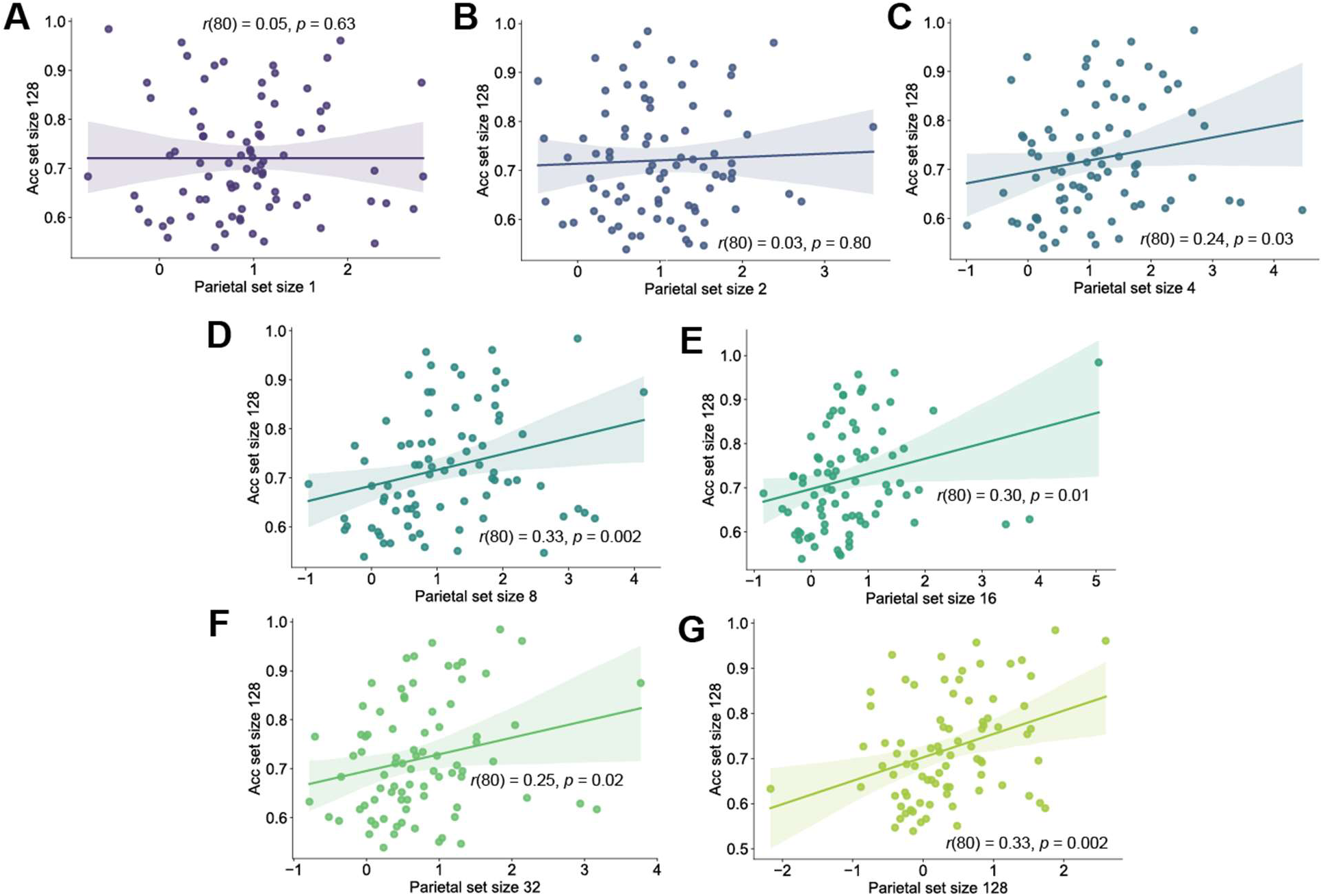
Parietal old/new effect for larger set sizes selectively predict long-term memory performance.

Taken together, these findings indicate that the parietal old/new effect selectively tracks individual differences in memory performance in a load-dependent manner: activity elicited by shorter lists primarily predicts WM ability, whereas activity elicited by longer lists more strongly predicts LTM ability. This pattern further supports the view that a common retrieval-related neural signal underlies both systems, while its behavioral relevance shifts as a function of memory load, consistent with our **convergence** hypothesis.

## Discussion

The present study examined whether recognition memory relies on qualitatively distinct neural mechanisms at different memory scales by tracking ERP signatures of old/new judgments across set sizes ranging from 1 to 128 items. Rather than revealing a sharp neural boundary corresponding to smaller memory scale within traditional working-memory (WM) capacity limits, the results demonstrated substantial continuity in recognition-related neural activity across memory scales. Specifically, both ERP and multivariate decoding analyses indicated systematic changes in the timing and strength of recognition-related neural signals as memory load increased. Both frontal and parietal old/new effects decreased in amplitude as list length increased, while their latencies became progressively delayed. Thus, increasing memory scales appears to alter the efficiency and timing of mnemonic processing rather than producing an abrupt transition to an entirely different neural recognition system.

These findings are particularly informative when considered in relation to the established ERP literature on recollection and familiarity. Recognition memory has traditionally been described as involving two forms of mnemonic information: familiarity, reflecting a sense that an item has been encountered previously without retrieval of specific contextual details, and recollection, involving retrieval of contextual or episodic information about the prior encounter (Yonelinas, 2001). ERP studies have identified an earlier mid-frontal old/new effect, often referred to as the FN400, that has been associated with familiarity, and a later parietal old/new effect or late positive component (LPC) that has been associated more strongly with recollection(Curran & Cleary, 2003; Rugg & Curran, 2007). Importantly, electrophysiological studies have provided evidence that these effects can be dissociated: the earlier frontal effect varies with familiarity strength, whereas the later parietal positivity is enhanced when information is recollected (Woodruff et al., 2006). The persistence of old/new effects across the present range of memory loads therefore suggests that recognition-related neural activity remains organized around processes that are observable across both relatively small and very large mnemonic sets.

At the same time, the present findings should not be interpreted as demonstrating that WM and long-term memory (LTM) use exactly the same retrieval mechanism. The systematic delay of recognition-related ERP effects at larger loads suggests that memory load changes when or how efficiently mnemonic evidence becomes available. This interpretation is compatible with the idea that increasing the number of studied items makes the relevant memory evidence less immediately accessible, resulting in slower accumulation or evaluation of evidence. However, the existence of similar ERP signatures at different loads does not establish that the same upstream retrieval operations generated those signals. Instead, qualitatively different retrieval processes could potentially converge onto a common downstream stage at which mnemonic evidence is evaluated and translated into an old/new decision. The distinction is important because ERP components are not perfectly process-specific.

Although the FN400 and LPC have traditionally been interpreted as neural correlates of familiarity and recollection, respectively, debate remains regarding the precise cognitive processes reflected by these components (Bridger et al., 2012; Rugg & Curran, 2007).

Consequently, the current findings provide stronger evidence for continuity in the neural architecture of recognition than for a fully shared retrieval mechanism.

The relationship between the present findings and the recollection/familiarity literature may therefore be better understood in terms of a common decision architecture operating on mnemonic evidence that can vary in strength and accessibility. Classic ERP research demonstrates that recognition decisions can be accompanied by temporally distinct neural signatures associated with familiarity and recollection (Curran & Cleary, 2003; Woodruff et al., 2006). The present results extend this framework by showing that recognition-related neural activity remains detectable even when the number of potentially remembered items increases far beyond conventional WM capacity. Rather than disappearing once memory load exceeds a small-capacity limit, the old/new effects become progressively weaker and later. This pattern is more consistent with a continuous scaling of mnemonic accessibility than with a categorical transition between WM and LTM retrieval systems.

The individual differences findings provide an important qualification to this interpretation. Although recognition may be supported by a common neural architecture across memory loads, the cognitive abilities that constrain performance need not remain constant. Recent behavioral evidence demonstrates a robust relationship between individual differences in WM capacity and visual LTM recognition, although the strength of this relationship depends on encoding conditions. Across four experiments involving 841 participants, Zhao and Vogel (Zhao & Vogel, 2025) found that WM capacity and attentional control predicted visual recognition memory as well as source memory in most conditions. However, this relationship was substantially reduced when items were presented rapidly enough to exceed attentional and WM limits during encoding. These findings indicate that WM ability can contribute to LTM recognition without implying that WM and LTM are the same memory system.

Other individual differences research similarly demonstrates a meaningful relationship between WM capacity and LTM performance. Miller et al. (Miller et al., 2019), for example, found that WM capacity was related to the intensity of attention during encoding and that individual differences in attentional engagement partially accounted for the relationship between WM capacity and LTM. Likewise, Unsworth et al. (Unsworth et al., 2014) found that individual differences in WM capacity were related to differences in access to information stored in LTM, suggesting that individual variability in memory performance can arise partly from differences in the accessibility of stored information. More recent modeling work further suggests that the relationship between WM capacity and LTM may reflect differences in both storage and retrieval processes, with evidence that high-WM individuals may be particularly effective at using associative encoding strategies (Streitberger et al., 2024). Together, these findings support the interpretation that WM and LTM are behaviorally interconnected even when their contributions to performance vary as a function of task demands.

Neural evidence also supports the possibility that a common recognition architecture can coexist with distinct sources of individual variability. Zhao et al. ((Zhao et al., 2026) recently demonstrated that EEG inter-electrode correlation patterns predicted individual differences in LTM encoding across set sizes of 32 and 128 items. Importantly, these neural predictions remained significant after controlling for visual WM and attentional-control performance, suggesting that LTM can possess a neural signature that is not reducible to general WM ability. Thus, a common neural format for recognition across memory loads should not be interpreted as evidence that the same cognitive ability determines successful performance at every scale. Instead, the neural system may preserve a common format for evaluating mnemonic evidence while the source of performance variability shifts depending on how much information must be maintained, encoded, or retrieved.

This distinction is particularly important for interpreting the present individual difference results. At smaller memory loads, successful recognition may benefit more strongly from the ability to maintain and manipulate a limited number of active representations, making WM capacity a relatively important predictor of performance. At larger loads, however, the number of representations exceeds conventional WM capacity, increasing reliance on durable LTM representations and the accessibility of those representations at retrieval. The fact that LTM ability becomes relatively more important at larger loads therefore does not necessarily contradict the neural continuity observed in the ERP data. Instead, the two findings may reflect different levels of analysis: the ERP results concern the neural architecture through which mnemonic evidence is evaluated, whereas individual differences results concern the cognitive resources and representations that determine how much usable evidence is available to that architecture.

## Conclusions

The present findings reveal substantial neural continuity in recognition memory across a broad range of memory demands. Rather than supporting a sharp distinction with increasing memory scales during recognition, the results suggest that recognition decisions are implemented through a common neural architecture whose timing varies as a function of memory accessibility. These findings are inconsistent with a strict **separate-retrieval** account and provide strong evidence for **convergence** with increasing memory scales during recognition. Importantly, however, the observed convergence does not support the strongest form of a **shared-access** account, which predicts that neural activity supporting retrieval with increasing memory scales should be mutually predictive of individual differences across individuals. Instead, the present results are consistent with a model in which small and large memory scales share aspects of the neural architecture supporting recognition while retaining partially distinct neural signatures. Thus, convergence between recognition memory retrieval with small and large memory scales need not imply that the two memory systems are fully interchangeable or that individual differences in their neural activity are necessarily shared. More broadly, these findings indicate that when memory is used to determine whether an item has been encountered before, the neural processes supporting recognition are substantially more continuous across memory scales than traditionally assumed, while still preserving meaningful distinctions between WM and LTM.

